# The hidden spatial dimension of alpha: 10 Hz perceptual echoes propagate as periodic travelling waves in the human brain

**DOI:** 10.1101/190595

**Authors:** Diego Lozano-Soldevilla, Rufin VanRullen

**Author notes:** **Corresponding author:** Rufin VanRullen, Centre de Recherche Cerveau et Cognition (CerCo), Pavillon Baudot CHU Purpan, BP 25202, 31052 Toulouse Cedex (France).

## Abstract

Alpha oscillations play a special role in vision. During sensory processing, reverse-correlation techniques revealed that white-noise luminance sequences elicit a robust occipital ∼10 Hz response that periodically reverberates the input sequence for up to 1 s. These perceptual echoes constitute the impulse response function of the visual system. However, the spatial dimension of perceptual echoes remains unknown: do they reverberate across the cortex simultaneously? Does stimulation over multiple visual coordinates evoke multiple synchronized echoes, or do they show consistent phase differences? Here, we tested the spatial dimension of perceptual echoes in two electroencephalogram (EEG) experiments manipulating the location of the visual stimulation. When a single disc flickered a white-noise luminance sequence in the upper visual field, we observed a single “echo wave” originating in posterior sensors and spatially propagating towards frontal ones (i.e. periodic travelling wave). The presentation of two independent flickering discs in separate visual hemifields produced two simultaneous and superimposed echo waves propagating in opposite directions, one in response to each stimulus. Strikingly, at many electrode sites, the phase of the two echoes differed, with a phase advance for the contralateral stimulus location. EEG source reconstruction tentatively located the waves within contralateral parieto-occipital cortex. In conclusion, the alpha rhythm processes stimulus information as a travelling wave that propagates across the cortical representation of retinotopic space in the human brain. In line with the “cortical scanning” hypothesis (Pitts & McCulloch, 1947), these results suggest the existence of an additional spatial dimension embedded in the phase of the alpha rhythm.

**Significance statement:** How does the spatial dimension of sensory processing relate to the temporal dimension of brain rhythms? Using correlation techniques, we characterized perceptual echoes, the average electroencephalogram response induced by visual stimuli that change luminance randomly. We found that perceptual echoes are actually periodic waves that travel through human visual cortex. Strikingly these periodic waves show consistent phase differences across the visual field, processing screen locations sequentially across distinct phases of the cycle following basic retinotopy. These results suggest the existence of an additional “hidden” spatial dimension in sensory cortex, encoded in the phase of the alpha oscillatory cycle. This could mean that perceptual echoes behave like sweeps of a sonar, processing the visual field in cycles of ∼100 ms duration.

## Introduction

The last decades have provided important insights regarding the functional consequences of neuronal oscillations (Engel et al., 2001; Fries, 2005; Buzsaki, 2006; Akam and Kullmann, 2014). Even at the level of single neurons and local populations, ionic conductances and circuits favor resonance at particular rhythms (Llinas, 1988). It is well documented that neuronal oscillations can modulate firing rate probability and they can improve stimulus information processing (Schaefer et al., 2006; Cardin et al., 2009; Sohal et al., 2009; Siegle et al., 2014).

The alpha rhythm (8 – 12 Hz) is the most prominent spectral fingerprint in the human brain and its functional role is an active area of research. For very brief stimuli at threshold, there is compelling evidence showing that both ongoing pre-stimulus alpha amplitude and phase modulate behavioral detection (Nunn and Osselton, 1974; Linkenkaer-Hansen et al., 2004; VanRullen et al., 2011; Vanrullen and Dubois, 2011). After stimulus onset, alpha oscillations are still present and there is a well-characterized amplitude reduction (i.e. desynchronization) that takes several cycles to recover (Pfurtscheller et al., 1996; Pfurtscheller and Lopes da Silva, 1999; Makeig et al., 2002). Conversely, brief visual stimulation also produces strong time-locked event-related potentials (ERPs) that last 100 – 500 ms and that are also correlated with visual perception (Cigánek, 1969; Hillyard et al., 1998; Luck et al., 2000). A recent study from our lab revealed a new time-locked ∼10 Hz oscillatory component that could last more than 1 s and that was thus dubbed “perceptual echo” (VanRullen and Macdonald, 2012). Stimulating participants with white-noise luminance sequences (i.e. equal power at all temporal frequencies) produced frequency-specific ∼10 Hz reverberations in posterior sensors (Ilhan and VanRullen, 2012; VanRullen and Macdonald, 2012). These echoes could be revealed by averaging the single-trial cross-correlations between the electroencephalographic (EEG) recordings and the luminance values of the random sequences (Lalor et al., 2006; Ilhan and VanRullen, 2012; VanRullen and Macdonald, 2012), yielding an estimate of the impulse response function (IRF) (Marmarelis and Marmarelis, 1978). In other words, the IRF is similar to an ERP that reflects changes in EEG brain activity proportional to luminance changes for different lags (i.e. cross-correlation). The perceptual echoes are a true response to the visual stimulus sequence: shuffling the pairing between luminance sequences and EEG time series practically abolishes the ∼10 Hz oscillation (VanRullen and Macdonald, 2012). The presence of long-lasting reverberations in this IRF (or perceptual echoes) implies that alpha oscillations are the optimal resonant frequency of the visual system, and that the brain can carry information about the external stimulation for more than 1 s (Ilhan and VanRullen, 2012; VanRullen and Macdonald, 2012; VanRullen et al., 2014; VanRullen, 2016).

It is not known, however, whether ∼10 Hz perceptual echoes from a particular visual field location reverberate across the cortex simultaneously or asynchronously. Moreover, does stimulation over multiple visual coordinates evoke multiple echoes, and are these oscillations uniform or do they show consistent phase differences? In order to investigate the spatial dimension of perceptual echoes, we re-analyzed the original dataset (VanRullen and Macdonald, 2012) to probe the existence of systematic phase variations across visual and/or cortical space. Here we demonstrate the hidden dimension of perceptual echoes: its spatial propagation. We show that perceptual echoes are periodic travelling waves that not only propagate across the cortex during several cycles but they also display phase differences between different spatial coordinates of the visual field, like the sweeps of a sonar.

## Materials and methods

A total of sixteen observers (n = 6, Experiment 1; n = 10, Experiment 2) participated in this study. All participants gave written informed consent before the experiment. The study was approved by the local ethics committee and was carried out in accordance with the provisions of the World Medical Association Declaration of Helsinki. Although findings from the same dataset have been published elsewhere (VanRullen and Macdonald, 2012), our new analysis with a focus on the unexplored spatio-temporal dimension of perceptual echoes reveals substantial new results. Electrophysiological activity was continuously acquired at 1024 Hz using a 64 channel ActiveTwo Biosemi EEG system. Horizontal and vertical eye movements were recorded using additional electrodes below the left eye and at bilateral outer canthi.

### Apparatus and Stimuli

Stimuli were generated using the MATLAB and Psychophysics toolbox (Brainard, 1997; Pelli, 1997; Kleiner et al., 2007) and displayed at 57 cm distance using a desktop computer (2.09 GHz Intel processor, Windows XP) with a cathode ray monitor (resolution: 640 × 480 pixels; refresh rate: 160 Hz) on a black background (luminance: 0.42 ± 0.02 cd/m2). Head position was stabilized using a chin rest fixed to the table. The display was calibrated and gamma corrected using a linearized lookup table.

The generation of visual stimuli was performed as follows. White-noise visual luminance sequences were displayed within a disc of 3.5 ° radius on a black background. In Experiment 1, a single disc was presented in the vertical meridian centered at 7.5 ° above the fovea. In Experiment 2, two independent white-noise luminance sequences were simultaneously displayed in two discs located in the left and right visual hemifields centered at 7.5 ° eccentricity on the horizontal meridian. In both experiments, the power spectrum of each randomly generated luminance sequence was normalized to have equal power at all frequencies. Each trial (6.25 s long) was initialized with random values from 0 to 1 drawn from a uniform distribution. Subsequently, we performed a Fourier transform of the time series, divided each resulting complex coefficient by its amplitude, and then applied an inverse Fourier transform to return to the time domain. The resulting time series were scaled to range from black (0.1 cd/m^2^) to white (59 cd/m^2^).

### Experimental design

A prototypical trial started with a white fixation dot presented at the center of the screen that remained throughout the experiment. Participants were told to keep fixation on the dot and covertly monitor the disc to detect a 1 s target square (3.75 °) appearing inside the disc on a random 25 % of trials. The target onset occurred at a random time within the sequence (uniform distribution, excluding the first and last 0.25 s). The area within the square followed the same sequence of luminance changes as the disc stimulus, but scaled in amplitude (using a staircase procedure) so that detection performance was fixed at approximately 82 %. Participants were instructed to press a button at the end of the sequence if they had detected the target.

### Analysis

EEG data were analyzed offline using FieldTrip (Oostenveld et al., 2011) (http://www.fieldtriptoolbox.org/); an open-source toolbox developed at the Donders Institute for Brain, Cognition, and Behavior (Nijmegen, The Netherlands) and custom Matlab code. The EEG data was re-referenced to the common average and down-sampled to 160 Hz. Trials of 6.25 s after the onset of the presentation of the luminance sequence were extracted. Blinks and electrocardiogram activity were isolated and removed by independent component analysis (Jung et al., 2000). Single trials of all participants were visually inspected and those containing excessive number of blinks and/or muscle contamination were rejected. In Experiment 2, at the beginning of the block, written instructions indicated to the participant whether they should pay attention to the left disc, to the right disc, or to both (3 block types). An exploratory analysis did not yield significant phase differences between attention conditions, and we thus pooled the data across the 3 block types to gain statistical power. Both target-present and target-absent trials were included in the cross-correlation analysis. This yielded a mean of 135.50 ± 24.24 and 463.90 ± 85.98 trials per participant in Experiments 1 and 2 respectively.

To obtain the IRF of the EEG we averaged the single-trial cross-correlations (Lalor et al., 2006; Ilhan and VanRullen, 2012; VanRullen and Macdonald, 2012) between the luminance sequence and the simultaneously acquired EEG time series at all lags between -0.2 to 0.7 s. Before cross-correlation analysis the first and the last 0.5 s of the two time series were excluded to avoid on/off transients (0.5 – 5.75 s).

For each electrode, phase differences were computed between perceptual echoes in response to left and right stimuli, and between electrodes for a given visual stimulation condition. For each participant, electrode and stimulation condition we band-pass filtered (8 – 12 Hz) the perceptual echoes using a 4^th^ order Butterworth filter. Subsequently, instantaneous analytic phase was obtained by taking the angle of the Hilbert-transformed band-pass filtered signal.

Template travelling wave propagation was computed by measuring all pair-wise phase differences *ψ*_*i*_ between the reference sensor POz and the rest of the EEG sensors. The template was based on the complex mean of phase differences over the significant time-samples (see below). An idealized wave template *W* was generated using 64 sine functions, each comprising the phase lag/lead *ψ* at sensor *i*:

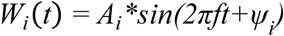

*f* was set at 10 Hz. The template was weighted by the amplitude of the corresponding perceptual echo peak amplitude *A*_*i*_ for sensor *i*.

### Dipole modelling

The wave templates were used to compute a source reconstruction of electroencephalogram (EEG) activity by equivalent current dipoles (ECD). A single equivalent dipole was fitted for each of 12 time points across a cycle of the wave template; we were interested to see how the 12 dipoles would change location and/or orientation across the cycle. Dipole fitting assumes that voltage activity measured at the scalp can be mathematically described using a small number of point-like ECDs (Scherg, 1990). First, standard electrode locations were co-registered to an anatomical MRI template (Holmes et al., 1998). This MRI was used to build a realistic shaped volume conduction model by means of the boundary element method (Oostenveld et al., 2001) and discretized into a grid with a 1 cm resolution. To source model the template travelling wave propagation, single dipole fitting was performed using nonlinear search to find, for every time point, the optimal dipole such that its location and moment (i.e. dipole orientation) minimize the difference between the model and the measured topography in the least squares sense (maximum iterations = 1000). More formally, let the signal be represented by the time series *X(t)* = [*x*_1_, *x*_2_, …, *x_N_*]^*T*^ where *N* is the number of time points reflecting the time varying cross-correlation values of the template echo function:

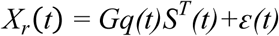

where *G* is the lead field matrix (channels by grid points) that weights how a given source *S* located in a specific grid position projects to the scalp sensors, and *ε* is the error term. The lead field matrix *G* is determined from the volume conduction model. The single dipole represents a location in Cartesian coordinates *q(t) = {q*_*x*_*, q*_*y*_*, q*_*z*_*}* with different magnitudes *||q(t)||* and moments *Θ(t) = q(t)/||q(t)|*| as a function of time. The goal of dipole fitting is to find, in our case, a single dipole whose magnitude and moment generate a source that minimizes the squared error *J*_*LS*_ between our data *X(t)* and the model:

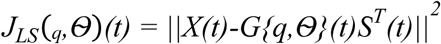

The optimal solution to minimize the least squares error at time point *t* is:

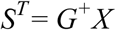

where *G{q,Θ}*^*+*^ is the pseudoinverse of *G{q,Θ}*. This results in the following expression:

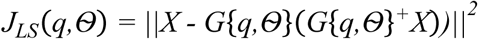

Dipole modeling of travelling wave propagation was performed on the template *W*_*i*_(*t*). The electric current dipole is an idealized model that is defined by two charges with equal and opposite polarity (Hämäläinen et al., 1993). This implies that a given source located in a specific brain area (i.e. fixed dipole location *q*), can project to the scalp either a positive or a negative voltage, only by rotating its moment by π radians. Due to the symmetry of our wave templates, the propagation of the negative trough of the template must therefore follow the same trajectory as its positive peak, but half a cycle later in time and with its dipole orientation inverted. To overcome this potential limitation, we also dipole-fitted the positive rectification of the wave template *W*_*i*_(*t*) as in (Hindriks et al., 2014), and we limited our fitting to one half of the wave cycle. To estimate the validity of our assumptions, we simulated three types of travelling waves and we reconstructed their scalp projections. To achieve this, we constructed a travelling wave model by setting the position and moment of a set of dipoles undergoing oscillatory activity fluctuations with systematic phase differences. We used the forward model *G* to estimate the potential field distribution *W* at the scalp level:

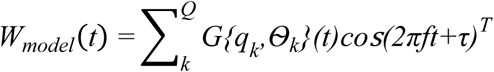

A global travelling wave was defined with dipoles (*Q* = 12) having different equally distant positions from occipital to frontal cortex, fitting a parabola function between the following three voxels expressed in MNI Cartesian coordinates (voxel 1 = [2.5cm -9.4cm -0.3cm], voxel 2 = [2.5cm -2.8cm 6.8cm], voxel = [2.5cm 2.5cm 5.3cm]; see Figure 4). Dipole moment was rotated linearly to cover π radians in 12 steps. The spanning was defined in spherical coordinates from π/3 to 4π/3 elevation (*φ*) while azimuth (*θ*) was locked to π/2 and transformed back to Cartesian coordinates. The parameter τ is what determines the specific phase-delay activation between dipole locations (*τ = 2π/Q*).

A Local periodic travelling wave was modelled with the same parameters as the Global travelling wave, but with the dipoles fixed to a single position, i.e. *q*_*k*_ *= q*_*1*_ for all *k*. The position was fixed to *q*_*1*_ = [2.5cm -7.1cm 4.4cm]. As for the global wave, the moment was rotated linearly to cover π radians in 12 steps from π/3 to 4π/3. The phase-delay activation between dipole locations was also *τ = 2π/Q*. This simulation also produced a posterior-to-anterior travelling wave on the scalp, but the underlying dipole rotation emulated a voltage propagation across local sulci/gyri (Hindriks et al., 2014).

The 2-dipole phase-lagged travelling wave was generated sitting two dipoles in occipital and parietal cortices (occipital: *q*_*1*_ = [2.5cm -8.8cm 1.5cm]; parietal: *q2* = [2.5cm -5.6cm 5.5cm]), with fixed moment (occipital: *φ* = 4*pi/3, θ = π/2; parietal: *φ* = 0.57*2*pi, θ = π/2). To simulate the activation lag between the dipoles, we set the parameter *τ = π/2*.

All models produce periodic travelling waves that propagate from posterior to anterior sensors (Hughes et al., 1992; Burkitt et al., 2000; Shevelev et al., 2000; Klimesch et al., 2007), similar to the one observed in Experiment 1.

### Statistical analysis

To estimate the time samples in which perceptual echoes in response to two simultaneous flickering sequences showed consistent phase relations across participants, we computed the phase-locking factor (PLV) across participants of the phase difference as follows:

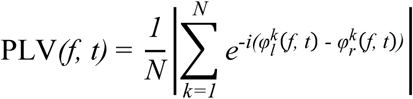

where *φ* is the alpha analytic phase of each stimulus condition (*l*, left; *r*, right), and *N* the number of participants. The PLV was averaged over sensors and compared to a surrogate distribution in which (i) the trials were randomly assigned to one of the two stimulus conditions, (ii) the PLV across participants was subsequently re-computed and combined across electrodes, and (iii) this procedure was repeated 6000 times. This way we determined the PLV distribution under the null hypothesis (i.e. no systematic phase relation across subjects between left vs. right stimuli). Statistical significance (p-values) was obtained by comparing the original PLV with the surrogate distribution for each respective time point. The false discovery rate (FDR) method was used to correct for multiple comparisons to avoid type I errors (Benjamini and Hochberg, 1995).

## Results

### Experiment 1: single visual location

Observers (n = 6) were stimulated with white-noise luminance sequences of 6.25 s duration at a single, fixed location, while EEG recordings were simultaneously acquired. Luminance changes were displayed within a peripheral disc presented above the fovea and participants had to covertly attend to it while maintaining fixation (Figure 1A). Cross-correlating the flickering sequences and the simultaneously acquired EEG recordings (Lalor et al., 2006; Ilhan and VanRullen, 2012; VanRullen and Macdonald, 2012), robust correlation values oscillating around the alpha rhythm were found in the IRF (Figure 1B). Although they differed in phase and frequency among participants, perceptual echoes appeared mainly over parieto-occipital leads. We observed that perceptual echoes did not reverberate in instantaneous synchrony (i.e. zero-phase lag) across the scalp. Instead, there were consistent phase differences between sensors, forming a spatio-temporal pattern of narrow-band oscillatory activity that propagated gradually through space (i.e. across the scalp) during several cycles, in other words, a periodic travelling wave (Hughes, 1995; Ermentrout and Kleinfeld, 2001). Specifically, posterior electrodes (POz, blue trace Figure 1B) lead the periodic wave propagation of activity relative to frontal electrodes (Fz, red trace Figure 1B). To characterize the spatial propagation of perceptual echoes, we performed a simple analysis computing a wave template (see Materials and methods section). We band-pass filtered the alpha component (8 – 12 Hz) of the perceptual echoes and we computed all pair-wise phase differences between a reference sensor showing strong echoes (i.e. POz) and the rest of the scalp, averaged over lags from 0.2 s to 0.5 s. Figure 1C shows the wave template evolution over a single cycle (100 ms). Perceptual echoes behaved as periodic travelling waves, whose propagation initiated over occipital sensors and flowed towards frontal sensors during several oscillatory cycles (Movie 1). This means that information processing for these stimuli, indexed by cross-correlation values in the IRF, was tightly linked to an alpha wave that travelled rhythmically in a posterior-to-anterior direction.

**Figure 1.**
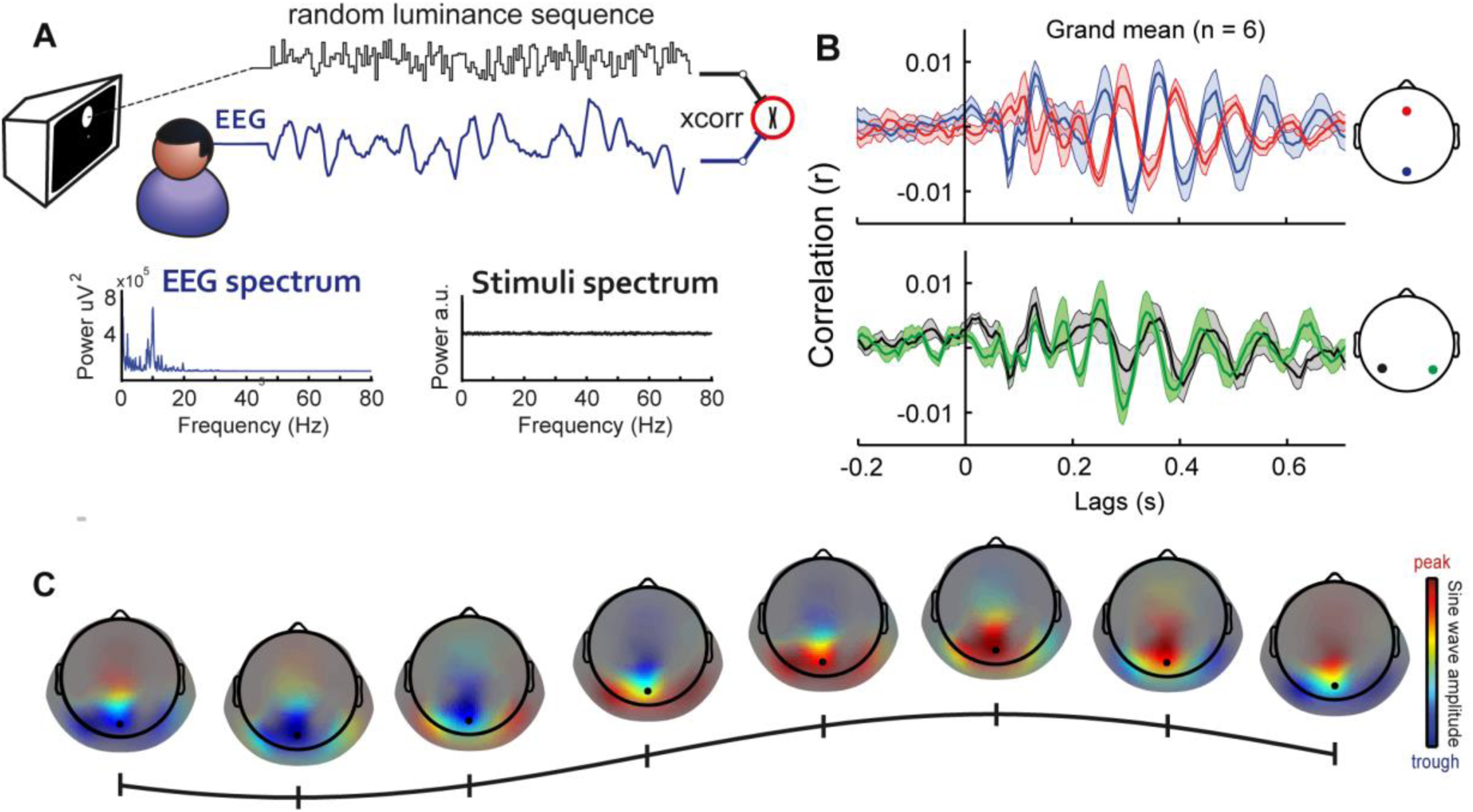
Visual input from a single location produces a bilateral, occipital-to-parietal oscillatory travelling wave. (A) In Experiment 1, random white-noise luminance sequences (6.25 s; black trace) were presented to the participants while EEG (blue trace) was simultaneously acquired. The sequences were displayed within a peripheral disc stimulus above the fixation point at a sampling rate of 160 Hz. To measure perceptual echoes the single-trial cross-correlations (Lalor et al., 2006; Ilhan and VanRullen, 2012; VanRullen and Macdonald, 2012) between the luminance sequence and the simultaneously acquired EEG time series were computed at all lags between -0.2 to 0.7 s and averaged (see Materials and methods section). Bottom, illustration of the power spectrum of a single-trial EEG and of a single-trial luminance sequence. (B) Grand-mean (n = 6) echo functions (shaded areas represent SEM across participants). While the perceptual echo in occipital sensors (e.g. POz in blue) showed a phase advance relative to frontal homologues (e.g. Fz in red) (top), perceptual echoes appeared synchronous between hemispheres (e.g. P5h, P6h shown in grey and green, respectively) (bottom). (C) Single-cycle wave template of the alpha component (8 – 12 Hz) of perceptual echoes. The template was computed by averaging over the time period 0.2 – 0.5 s the analytic pair-wise phase differences between a reference electrode (POz in black) and the rest of the EEG (see Materials and methods section). Color code represents the peak (trough) of the wave relative to the reference electrode. For visualization purposes, topographies were masked with grey transparency using alpha transparency values proportional to the peak amplitude of the perceptual echo at each electrode, raised to the power of three.

### Experiment2: two simultaneous visual locations

The above observations suggest that perceptual echoes do not emerge synchronously over the entire scalp. To go one step further, we analyzed the data of a second experimental condition, in which two independent white-noise luminance sequences were simultaneously presented on respective discs to the left and right of the fixation point (Figure 2A). Can a single cortical region (or scalp channel) simultaneously respond with an echo to each visual location? If yes, would the two echoes unfold synchronously, or show consistent phase differences? Experiment 2 (n = 10) allowed us to test two spatial dimensions of perceptual echoes: visual (i.e., comparing screen coordinates) and cortical (i.e., comparing scalp coordinates).

**Figure 2.**
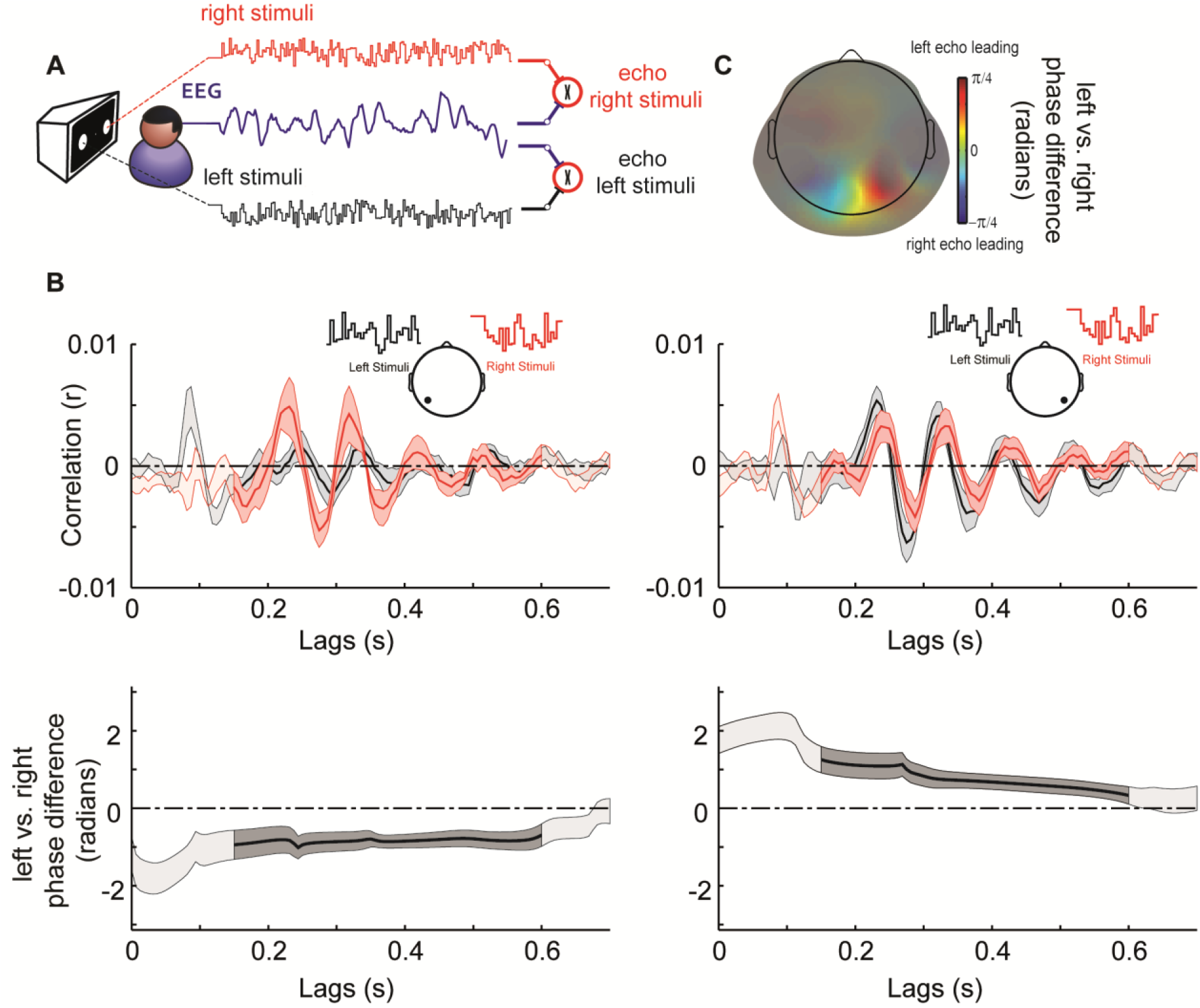
Spatial propagation of perceptual echoes in the visual domain (phase differences between screen locations). (A) In Experiment 2, two independent random white-noise luminance sequences (6.25 s long) were presented to the participants while EEG was simultaneously acquired. The sequences were displayed within two peripheral discs located left and right of the fixation point at a sampling rate of 160 Hz. Two sets of single-trial cross-correlations between each stimuli sequence and the EEG were carried out, to compute the average echo in response to right stimuli (in red throughout the figure) and the average echo in response to left stimuli (in black). (B) Top. Grand-mean (n = 10) perceptual echoes measured at a single sensor of interest (P5h on the left-hand side, P6h on the right-hand side) in response to the two stimuli sequences. Shaded areas represent SEM across participants and areas with high contrast represent significant phase relation consistent across participants (PLV) (0.15 – 0.6 s; see Materials and methods section). Topographic inserts represent the sensors of interest where perceptual echoes in response to the two independent luminance sequences were measured. Bottom. Time-resolved analytic phase differences (left minus right echoes) of the alpha band component (8 – 12 Hz) of the two perceptual echoes measured in one sensor of interest. Shaded areas represent SEM across participants and areas with high contrast represent significant phase relation consistent across participants (PLV). Here we see that the contralateral stimulus location had a systematic phase advance relative to the ipsilateral one (C) Grand-mean topographic representation of phase differences (left minus right perceptual echoes) averaged over the time period showing a significant phase relation consistent across participants (PLV; 0.15 – 0.6 s). Color code represents phase differences in radians. For visualization purposes, topographies were masked with grey transparency using alpha transparency values proportional to the peak amplitude of the perceptual echo at each electrode, raised to the power of three.

To test how perceptual echoes differed across distinct visual coordinates (i.e. stimuli located in the left and right visual hemifields), we computed cross-correlation analysis between the EEG and each luminance sequence separately (Figure 2A). On each occipito-parietal EEG sensor, we found two independent echo responses evoked by the respective stimuli sequences (Figure 2B). In order to estimate the time lags for which perceptual echoes in response to the two simultaneous flickering sequences showed a systematic phase relation, we computed the PLV across participants of the phase difference between the two echoes. Briefly, for each participant and EEG sensor we bandpass filtered the alpha component of perceptual echoes in response to the two flicker sequences. We obtained the analytic phase of the filtered alpha band using the Hilbert transform. Then, for each participant we computed the magnitude of the phase difference between the left and right perceptual echoes; the PLV for this phase difference was finally computed over participants (and averaged over all sensors, see Materials and methods section). We found that participants showed a consistent phase relationship between the echoes evoked by the discs located in each visual hemifield over the lag period from 0.15 s to 0.6 s (p < 0.01, comparison with surrogate distribution of PLV under the null hypothesis, FDR corrected). Note that these significant PLV values do not inform about the magnitude of the phase difference (only about the consistency of phase relations). For example, it could be the case that both echoes were always measured in each EEG electrode synchronously (i.e. zero phase-lag) or there might be a systematic lag/lead between responses (i.e. periodic travelling wave). To disentangle between the two possibilities, we examined the direction of the phase relationship. For this, we focused on sensors of interest located contralateral to the site of stimulation, and we computed the alpha band analytic phase differences between left v.s. right perceptual echoes (Figure 2B). While the left sensor revealed significant negative non-uniform phase differences (circular mean = -0.7743; Rayleigh z = 6.01; p = 0.0011), the right one showed complementary positive non-uniform phase differences (circular mean = 0.7104; Rayleigh z = 5.7616; p = 0.0015). These results indicate that perceptual echoes measured in contralateral sensors were leading (i.e. statistically significant phase advancement) relative to their ipsilateral homologues.

The cortical dimension of perceptual echoes refers to the periodic wave propagation across the scalp corresponding to visual information from a particular screen location. Does a given stimulus produce an echo in a single cortical source, resulting in instantaneous synchronization over electrodes, or do sensors show systematic phase-lags? In case of systematic phase differences, what is the propagation direction? This question was already addressed in Experiment 1 for a single-stimulus situation, but the observed periodic travelling wave may have distinct properties (or disappear altogether) when two stimulus locations must be simultaneously processed. As in Experiment 1, we mapped the analytic phase of the alpha component of perceptual echoes in response to each given stimulus, separately for left and right sensors. For the left stimulus, we found statistically significant negative phase differences, meaning that right parietal sensors showed a phase advance relative to the left sensors (Figure 3A). Complementary results (i.e. positive phase difference) were found for the right stimulus. In both analyses, the mean phase difference was around pi/3 corresponding approximately to a ∼16 ms delay (Left stimuli: left minus right sensor, circular mean = -1.0086 radians; Rayleigh z = 5.1873; p = 0.0033. Right stimuli: circular mean = 0.9615 radians; Rayleigh z = 3.8284; p = 0.0177). As done previously, we characterized the spatiotemporal propagation of perceptual echoes by computing a wave template. For each perceptual echo in response to each stimuli sequence, we computed all pair-wise phase differences between a reference sensor of interest (POz) and the rest of the EEG. We found that each template wave propagated mainly in the contra-to ipsi-lateral direction (Figure 3B). Strikingly, these two travelling waves happened simultaneously and were therefore superimposed on the scalp (Movie 2); they were disentangled here by reverse-correlation techniques.

**Figure 3.**
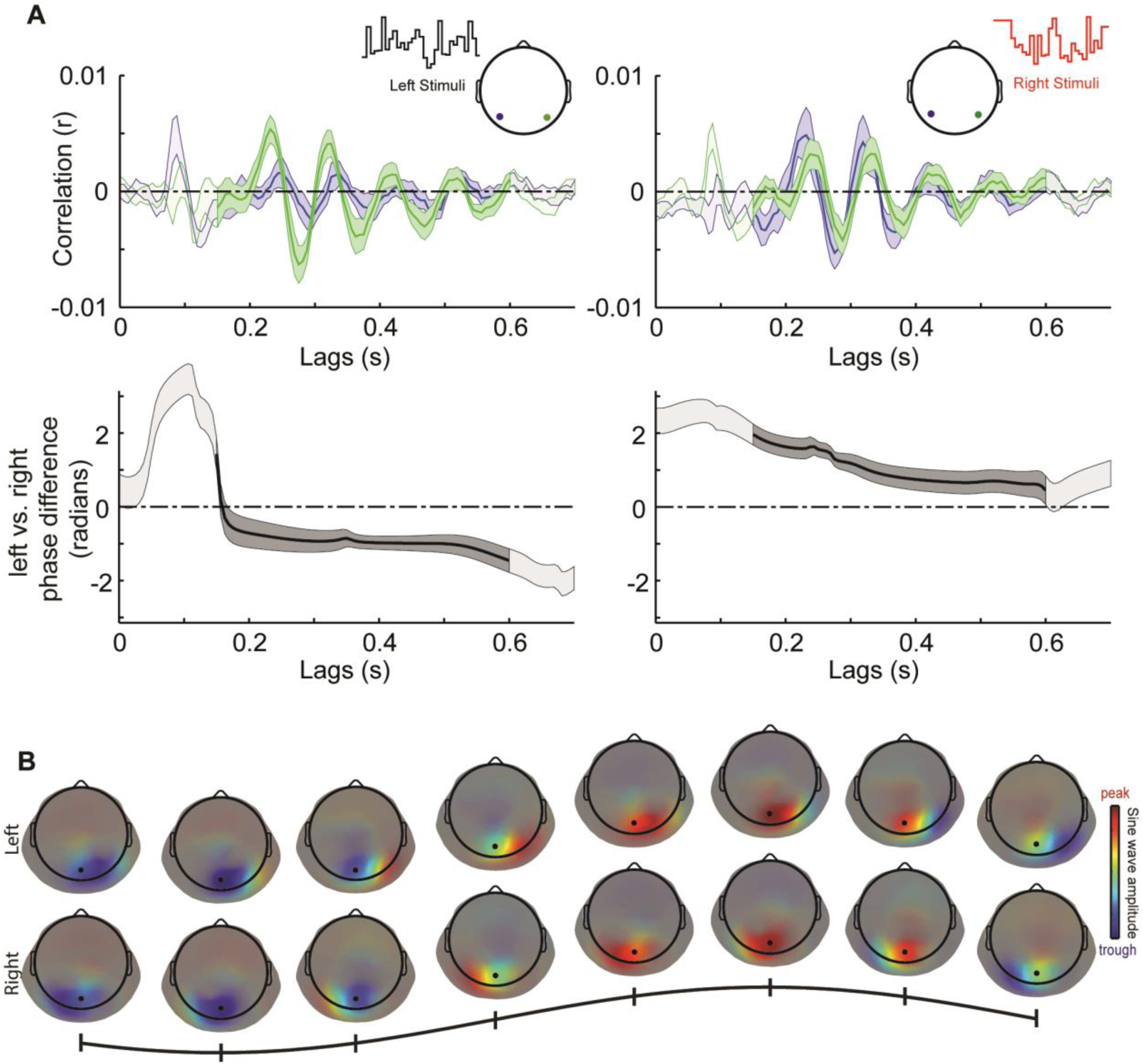
Spatial propagation of perceptual echoes in the scalp domain (phase differences between scalp locations). (A) Top. Grand-mean (n = 10) perceptual echoes measured at two sensors of interest (P5h in blue and P6h in green) in response to left stimuli sequences (left-hand side) or right stimuli sequences (right-hand side). Shaded areas represent SEM across participants, and areas with high contrast represent significant phase relation consistent across participants (PLV) (0.15 – 0.6 s; see Materials and methods section). Topographic inserts represent the sensors of interest. Bottom. Time-resolved analytic phase differences (left minus right electrodes) of the alpha band component (8 – 12 Hz) of the perceptual echoes. Shaded areas represent SEM across participants and areas with high contrast represent significant phase relation consistent across participants (PLV). Here we see that the contralateral scalp location had a systematic phase advance relative to the ipsilateral one (Rayleigh test, p<0.005 for left stimuli, p<0.02 for right stimuli). (B) Single-cycle wave template of the alpha component (8 – 12 Hz) of perceptual echoes in response to left (top) and right (bottom) luminance sequences. The template was computed by averaging over the time period 0.15 – 0.6 s the analytic pair-wise phase differences between a reference electrode (POz in black) and the rest of the EEG (see Materials and methods section). Color code represents the peak (trough) of the wave relative to the reference electrode. For visualization purposes, topographies were masked with grey transparency using alpha transparency values proportional to the peak amplitude of the perceptual echo at each electrode, raised to the power of three.

### Source modelling

Finally, we chose ECD modelling to tentatively reconstruct sources of the periodic travelling waves found on scalp EEG in the two experiments. Due to limitations of inverse modelling (Michel et al.; Lütkenhöner, 2003; Destexhe and Bedard, 2012; Riera et al., 2012; Gratiy et al., 2013) and the a priori unknown ground truth of the spatial configuration of the sources underlying periodic travelling waves, we dipole-fitted both the original (unrectified) wave template and its positive-rectified version. We built simulated datasets to demonstrate the rationale behind this approach. Using a realistic biophysical forward modelling, we generated three types of periodic travelling waves: Global (12 anatomically separated dipoles simultaneously oscillating but with a systematic phase lag), Local (12 anatomically co-localized dipoles of different orientations, simultaneously oscillating but with a systematic phase lag) and the 2-dipole phase-lagged (with only 2 oscillating sources having a phase-lag of π/2; see Materials and methods section). All three situations could produce travelling waves qualitatively comparable to the ones observed experimentally (Figure 4B). Fitting single ECD for each time point to both the positive-rectified template and the unrectified template, we empirically showed that positive rectification provides a more accurate description when the ground truth is a Global periodic travelling wave, whereas dipole position and orientation can be recovered more accurately using the unrectified template when the ground truth is a Local or a 2-dipole phase-lagged periodic travelling wave (Figure 4). Given that we do not know the ground truth for our empirical data, we reported the two solutions.

**Figure 4.**
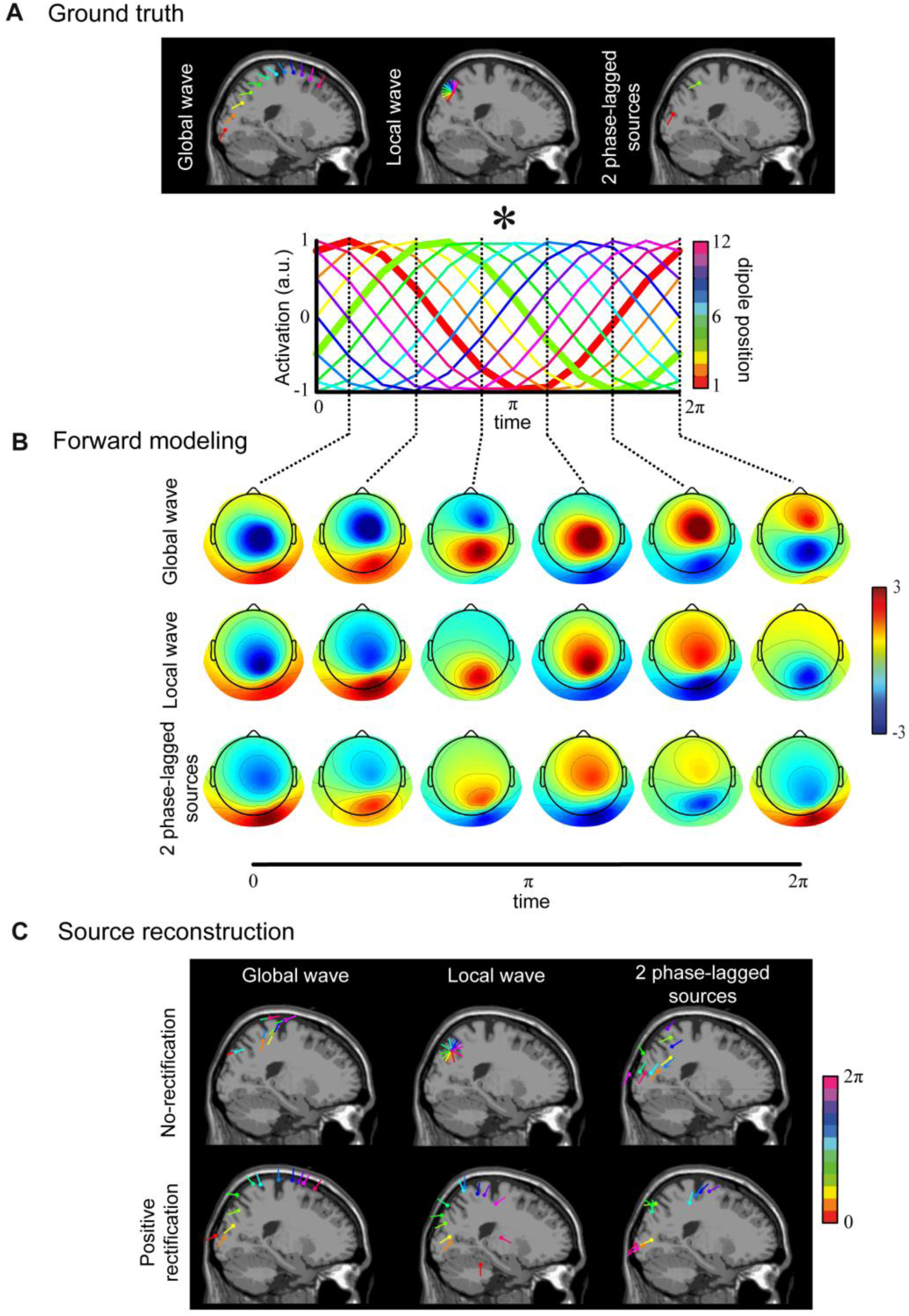
Equivalent current dipole (ECD) source reconstruction on simulated model travelling waves. (A)Top: Ground truth dipole positions used to simulate a Global (left), Local (middle) and 2-phase-lagged (right) travelling waves. Bottom: activation cosine functions used to simulate travelling waves over one cycle (100 ms). The travelling wave is modelled using either 12 (Global and Local travelling waves) or 2 (2-dipole phase-lagged travelling wave) concurrent dipoles whose activations across time are governed by a cosine function with consistent phase differences. Color code represents dipole position. The Global wave is produced by spatially separated dipoles, whereas the Local wave originates from overlapping dipoles with different spatial orientations. The 2 phase-lagged travelling wave is produced by two separated sources with a fixed phase-lag between them of π/2 (bold red and green activation functions). (B) Forward modelling was applied to the three simulated travelling waves (ground truth). Six topographic representations of successive time samples for Global (top), Local (middle) and 2-dipole phase-lagged (bottom) travelling waves illustrate the propagation of the wave on the scalp. Color code represents the strength of field distribution (arbitrary units). (C) Source reconstruction of the scalp projection of simulated travelling waves illustrated in B. Single equivalent current dipole modeling of simulated Global (left), Local (middle) and 2 phase-lagged (bottom) travelling waves was applied on un-rectified (top, No-rectification) and rectified topographical data (bottom, Positive rectification) at 12 distinct time points within a cycle. Dipole length represents the wellness of fit for the corresponding time point. Color code represents the 12 time samples that covered a single cycle of a simulated travelling wave. In conclusion, the positive rectification method provides the most accurate fitting results when the scalp pattern (in B) is actually produced by a Global travelling wave in cortex (ground truth in A). In contrast, the non-rectification method provides the best fitting results when the scalp pattern (in B) is caused by either Local or 2-dipole phase-lagged travelling waves (ground truth in A). As the ground truth for our experiments is unknown, both methods must be used when fitting dipole sources to the scalp travelling wave.

In Experiment 1 (single disc above fovea), the tentative ECD source reconstruction based on the positive rectified template suggested that the underlying wave likely propagated over bilateral parieto-occipital regions (occipital superior lobe, cuneus, precuneus and parietal superior lobe; Figure 5, upper row). The non-rectified wave template yielded a more spatially constrained solution centered around bilateral parietal lobe (precuneus; Figure 5, lower row). Concerning Experiment 2 (two discs located in left and right visual hemifields), ECD analysis on the positive rectified wave template tentatively revealed that each of the two waves propagated within the contralateral hemisphere relative to the stimulation site, from occipital to parietal cortex (mid temporal lobe, mid and superior occipital lobe, cuneus and precuneus, inferior and superior parietal lobe until the post-central midline; Figure 6). As in Experiment 1, the source localization of the non-rectified wave template produced clustered dipoles in regions of parieto-occipital lobe (mid and superior occipital lobe, cuneus and precuneus, inferior and superior parietal lobe; Figure 6). In conclusion, both ECD solutions (based on rectified or non-rectified templates) agreed on the likely parieto-occipital origin of the periodic travelling waves; however, the exact spatial extent of these waves was considerably larger for rectified than non-rectified templates, and thus remains difficult to determine.

**Figure 5.**
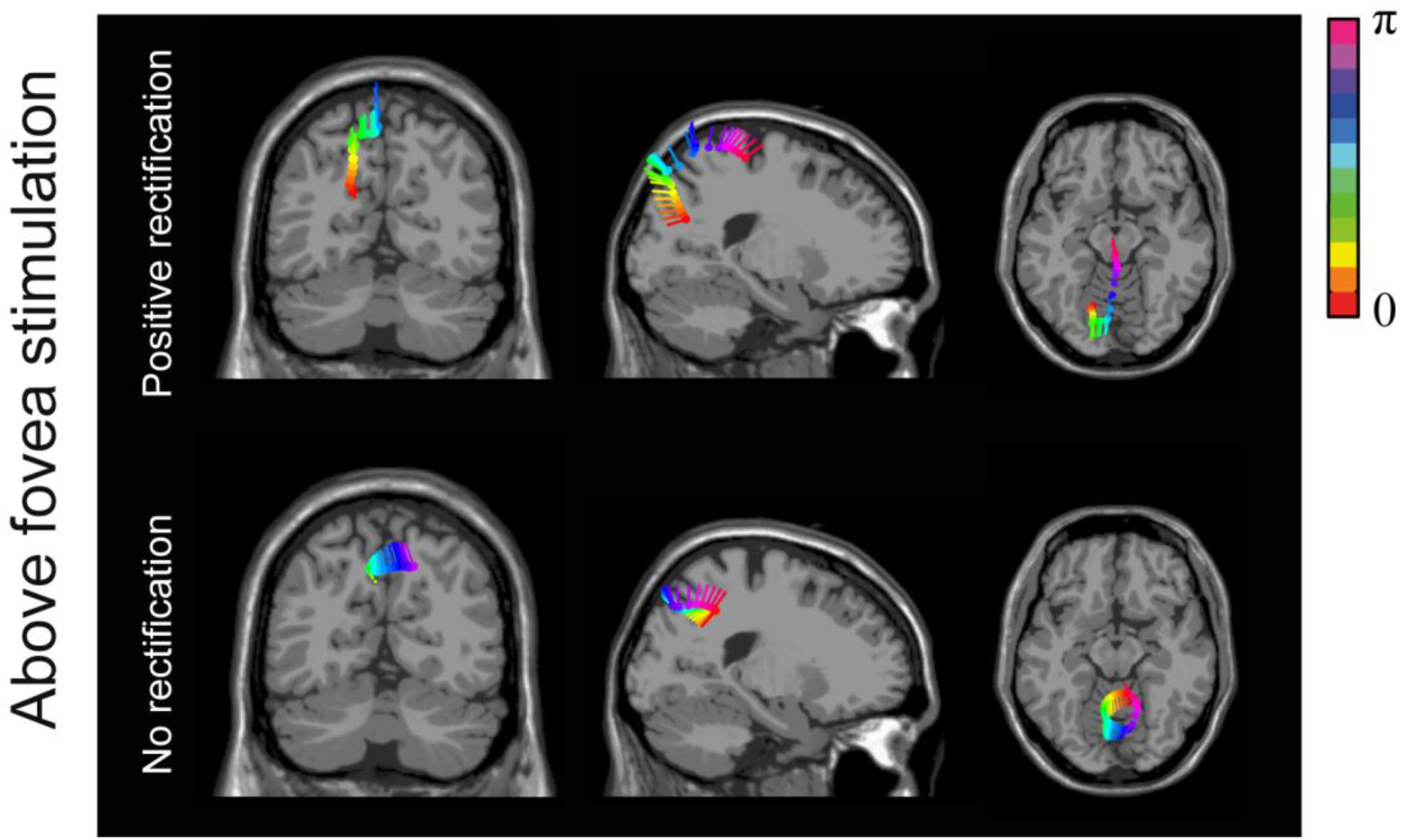
Equivalent current dipole (ECD) analysis of perceptual echoes from Experiment 1. Horizontal, sagittal and coronal planes of dipole positions of the single-cycle wave template obtained in Figure 1C. Size of each dipole represents wellness of fit for the fitted time point, which ranged from 74 % – 95 %; dipole color represents the phase of the wave at electrode POz at the fitted time point (see Materials and methods section). Both the positive-rectified and the unrectified fitting methods place the wave in occipito-parietal regions. As expected (see Figure 4), the travelling wave appears to propagate over a larger cortical extent using the positive-rectified method compared to the unrectified method.

**Figure 6.**
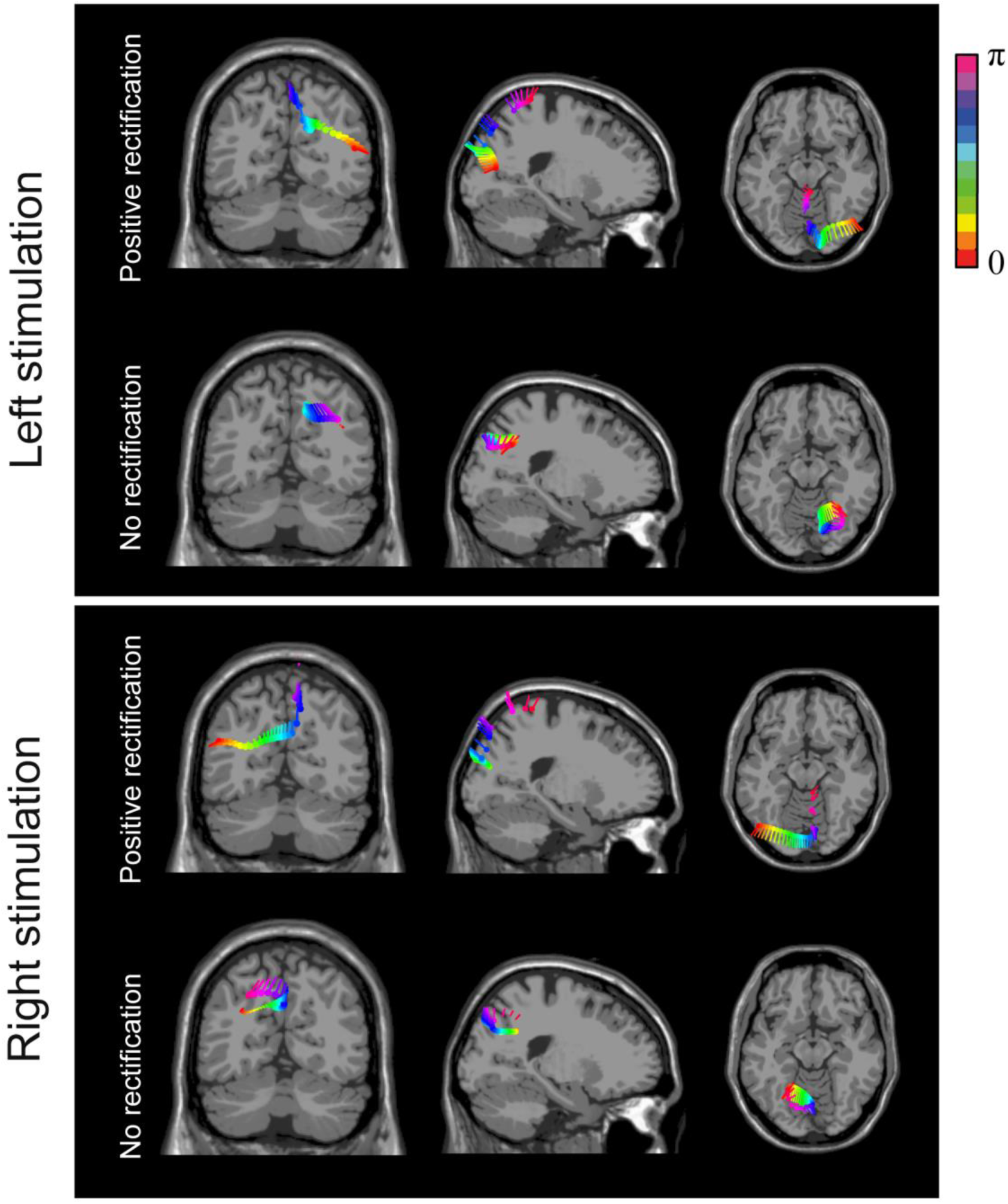
Equivalent current dipole (ECD) analysis of perceptual echoes from Experiment 2. Horizontal, sagittal and coronal planes of dipole positions of the single-cycle wave templates obtained in Figure 3B. Size of each dipole represents wellness of fit for the fitted time point, which ranged from 71 % – 93 %; dipole color represents the phase of the wave at electrode POz at the fitted time point (see Materials and methods section). The overall spatio-temporal patterns of dipoles are consistent with those observed in Experiment 1 (Figure 5), except that occipito-parietal regions here are systematically activated in the hemisphere contralateral to the stimulus location, in line with the retinotopic organization of visual cortex. As previously (Figures 4-6), the travelling wave appears to propagate over a larger cortical extent when using the positive-rectified method compared to the unrectified method.

## Discussion

In the two experiments presented here, we have tested the spatial dimension of perceptual echoes by manipulating the location of visual stimulation and measuring the EEG responses across the scalp. In Experiment 1, the stimulus located above the fovea generated a single periodic travelling wave, to which the two hemispheres contributed with coherent activity, propagating from posterior to frontal sensors during multiple alpha cycles. In Experiment 2, we showed that two independent random flickering sequences located in the two visual hemifields generated two superimposed periodic travelling patterns, whose phase propagation travelled following the contra-to ipsi-lateral direction. As a result, the two perceptual echoes recorded at a given scalp location showed a sequential activation: periodic waves in response to contralateral stimulation were phase-advanced relative to ipsilateral stimulation.

We can be reasonably confident that the periodic travelling waves recorded across the scalp surface in the two experiments actually reflect spatial propagation of activity within and/or between specific brain regions. Because electrical conduction delays are negligible, a single static source would necessarily produce synchronized oscillations across the entire scalp, with opposite polarities on the two sides of the equivalent current dipole, but no smooth phase transitions (this is typically called a “standing wave”). Thus, in order to produce smooth phase changes, there must be at least two or more underlying oscillatory sources with a consistent phase difference between them; this is actually the minimal definition for a “periodic travelling wave” (Hughes, 1995; Ermentrout and Kleinfeld, 2001). Nonetheless, we acknowledge that whether these multiple oscillatory sources span a large cortical extent or a very small one (making the effect mathematically similar to a rotating dipole) remains an open question; our ECD modelling efforts (Figures 4-6) suggest that both local and global hypotheses (as well as intermediate ones) are viable. With this in mind, the present results demonstrate that perceptual echoes behave like periodic travelling waves. To our knowledge, this is the first demonstration of two concurrent and overlapping stimulus-induced periodic travelling waves in humans.

While previous resting-state human EEG studies reported anterior to posterior propagation dynamics (Nunez et al., 2001; Nolte et al., 2008), visual stimulation experiments produced ERPs containing an alpha frequency component that propagated from posterior to anterior sensors (Hughes et al., 1992; Burkitt et al., 2000; Shevelev et al., 2000; Klimesch et al., 2007). The present results constitute a significant step forward for several reasons. First, contrary to ERPs that measure voltage variations and can only be indirectly related to stimulus properties (Fellinger et al., 2012; Alexander et al., 2013), the perceptual echoes (or so-called IRFs) directly reflect sensory information processing, indexed by cross-correlation coefficients between EEG and luminance sequences. Second, the resulting correlations (i.e. perceptual echoes) obeyed simple retinotopic rules (i.e. contra-to ipsi-lateral propagation), which implies that they follow the structure and functional organization of basic cortical pathways. Third, we were able to simultaneously induce two overlapping periodic travelling waves propagating in distinct directions (Movie 2). Finally, another aspect of the present travelling waves worth insisting on is their periodic nature. Classic studies in monkeys have found travelling waves mainly during resting-state conditions (Ermentrout and Kleinfeld, 2001), but more recent studies have shown that sensory stimulation using periodic stimuli also produces travelling waves, whereby for each new stimulus cycle a single wavefront propagates through cortical space (Benucci et al., 2007; Nauhaus et al., 2012; Sato et al., 2012; Muller et al., 2014). However these travelling waves are different from the periodic travelling waves found here in which perceptual echoes evoked by random (aperiodic) stimulation maintain their periodicity during four or more consecutive cycles.

What could be the neurophysiological mechanisms underlying the consistent phase differences we observed at the scalp level? A straightforward explanation could be that perceptual echoes propagate across a large portion of cortical space. More concretely, multiple neuronal patches of occipito-parietal cortex could be activated in a chain reaction (i.e. phase delays between neuronal pools); in this way, the scalp phase-differences would directly mirror underlying cortical phase differences. Our ECD analysis on the positive rectified topographical template suggests that this is indeed one valid possibility (Figures 4-6). However, due to the well-known limitations of noninvasive source reconstruction (Michel et al.; Lütkenhöner, 2003; Destexhe and Bedard, 2012; Riera et al., 2012; Gratiy et al., 2013), there are alternative explanations to consider. Mathematically, a periodic travelling wave measured at the scalp can also be reduced to a static dipole(s) whose orientation rotates as a function of time, producing an “apparent” propagation that can be measured with EEG by volume conduction (Nunez and Srinivasan, 2006). A hypothetical pool of neurons would synchronize its firing activity around the alpha band, and its dipole moment would rotate in space (e.g. from contra-to ipsi-lateral direction along the parieto-occipital cortex; Figure 4). This would produce consistent phase differences in the sensor domain, because the dipole would project its peak electrical voltage at various points in the channel space as a function of time. However, this mathematically valid construct has no direct physiological equivalent: taking the rotating dipole hypothesis literally would imply that the axons of the hypothetical pool of neurons would physically rotate around the soma every 100 ms! A more plausible physiological interpretation would consist in a spatially restricted wave activation pattern that propagates locally, through sulci and gyri of varying orientation, resulting in a global periodic travelling pattern at the scalp level. This is known as the intra-cortical hypothesis, which assumes that phase-differences at the scalp level reflect the geometric curvature of the wave propagation (Hindriks et al., 2014). In this case, however, there would still be a periodic wave of activity travelling over cortical space, only with a much more restricted spatial extent. Our ECD modelling on the unrectified wave template is also compatible with this interpretation. A third possibility, the 2-dipole phase-lagged travelling wave in which only 2 dipoles oscillating with a fixed phase difference can be thought of as an intermediate point in a continuum between the global and local situations. Future studies using magnetoencephalographic and preferably electrocorticographic (ECoG) recordings would be instrumental in determining the origin and propagation pattern of perceptual echoes. A preliminary report from human ECoG shows that alpha oscillations were present over multiple brain areas and they propagated in a posterior-to-anterior direction (Jacobs et al., 2017). Another recent ECoG study found that sleep spindle activity forms spontaneous circular travelling waves that repeat themselves multiple times over several hours and that might help to store and integrate memories in humans (Muller et al., 2016).

Overall, it is unavoidable that the periodic travelling wave observed at the scalp originates from a travelling wave in the brain, although we cannot precisely pinpoint its spatial extent: local propagation vs. large-scale propagation over multiple brain regions. Depending on its exact spatial extent, the spatial propagation of the travelling wave may reflect several (non-exclusive) underlying processes: (i) a (periodic) feed-forward pass of activity through the hierarchy of visual areas, from V1 to posterior parietal cortex, (ii) horizontal connections linking various portions of retinotopic space, potentially through iso-eccentric functional connections (Arcaro et al., 2011; Arcaro and Kastner, 2015; Arcaro et al., 2015), and/or (iii) inputs from subcortical structures of the thalamus (Gücer et al., 1978), such as the pulvinar (Albe-Fessard et al., 1966; Albe-Fessard, 1973; Lopes da Silva et al., 1973), a neural communication “hub” that has been demonstrated to participate in the oscillatory coordination of activity between occipital and parietal areas at alpha frequency (Saalmann et al., 2012) Future studies would be needed to disentangle these underlying neural mechanisms.

What could be the functional advantage of a travelling perceptual echo spanning several cycles? We showed consistent phase differences in the scalp domain (i.e. when comparing different sensors) and in the visual domain (i.e. when comparing the echoes produced by two simultaneous flickering stimuli located in separate hemifields). This could be explained as a wave (or a number of simultaneous waves) sequentially scanning different parts of the cortex as a function of visual stimulation coordinates. Taking an analogy, perceptual echoes would implement a sort of sonar sweep across the visual field. Every cycle of the wave would sweep the visual space from contra-to ipsi-lateral positions, sampling the visual world rhythmically and propagating it through the different parts of the cortex sequentially (Movie 3). This is precisely what Pitts & McCulloch conjectured 70 years ago in the "cortical scanning" hypothesis: *“…this alpha rhythm performs a temporal "scanning" of the cortex which thereby gains, at the cost of time, the equivalent of another spatial dimension in its neural manifold”* (Pitts and McCulloch, 1947). Here, we corroborated, for the first time, the existence of such an additional spatial dimension in sensory cortex, encoded in the phase of the alpha oscillatory cycle of perceptual echoes.

## Author Contributions

D.L-S. and R.V. wrote and reviewed the manuscript. D.L-S. analyzed data. R.V. designed and performed research.

## Acknowledgments

This work was supported by an ERC Consolidator grant P-CYCLES (nr: 614244) awarded to R.V. We thank James Macdonald for help with data collection.

